# Maternally-transferred thyroid hormones and life-history variation in birds

**DOI:** 10.1101/775981

**Authors:** Bin-Yan Hsu, Veli-Matti Pakanen, Winnie Boner, Tapio Eeva, Blandine Doligez, Ton G.G. Groothuis, Erkki Korpimäki, Toni Laaksonen, Asmoro Lelono, Pat Monaghan, Tom Sarraude, Barbara Tschirren, Robert L. Thomson, Jere Tolvanen, Rodrigo A. Vásquez, Suvi Ruuskanen

## Abstract

Life-history traits vary largely across species and several physiological parameters have been proposed to be associated with life-history variation, such as metabolic rates, glucocorticoids, and oxidative stress. Interestingly, the association between thyroid hormones (THs) and life history variation has never been considered, despite a close interaction between THs and these physiological traits. Because of the crucial effects on embryonic development, THs can also induce transgenerational plasticity when transferred to developing offspring, for instance, via egg yolks in birds. In this study, we compiled a unique data set of maternal yolk THs in 34 bird species across 17 families and 6 orders, and tested for associations with various life-history traits. Our phylogenetic mixed models indicated that both concentrations and total amounts of the two most important forms of THs (T3 and T4) were higher in the eggs of migratory species than in those of resident species, and that there were higher total amounts of T3 in the eggs of precocial species than in those of altricial species. However, maternal THs did not show clear associations with any traits of the pace-of-life syndrome, such as developmental duration, growth rate, or lifespan. When taking environmental factors into account, we found that captive species deposited higher TH concentrations and larger amounts in the egg yolks than wild species. These findings suggest that maternal THs are likely involved in the evolution of life-history variation, or vice versa.

## Introduction

Life-history variation constitutes a large part of biodiversity. Across species, life-history traits are often inter-correlated and present along several continuums. For example, many life-history traits, such as lifespan, fecundity and growth rates, co-vary with one another and constitute a continuum representing the ‘pace of life’ (Ricklefs and Wikelski 2002; Dammhahn et al. 2018). On the one end of this continuum, species exhibit fast growth and development, early maturation, but also fast senescence and short lifespan (“fast” species). On the other end, species have slow growth and development, late maturation and first reproduction, but also live longer (“slow” species). Other life-history continuums, such as the nutrient source for developing embryos (from yolk or via placenta), income versus capital breeders, precocial to altricial developmental modes, and various degrees of migratory tendencies, also exist across various taxa in the animal kingdom.

Identifying the mechanisms underlying the evolution of life history differences has fascinated evolutionary biologists for decades. For example, metabolic rate has been proposed as a mediator for the pace-of-life (POL) continuum, as it exhibits a parallel pattern with fast species usually having higher basal and resting metabolic rates (BMR and RMR), while slow species having lower metabolic rates (e.g. Wikelski et al. 2003; Auer et al. 2018). Some studies also proposed other possible mediators, such as glucocorticoids (Hau et al. 2010), oxidative stress (Vágási et al. 2019), and blood glucose level (Tomasek et al. 2019). We propose that thyroid hormones (THs), which have regulatory functions on metabolism (Mullur et al. 2014) and interactions with glucocorticoids (Darras et al. 1996; Kulkami and Buchholz 2012; Cortés et al. 2014), may be an additional mechanism underlying life-history variation, such as the POL continuum, or explain previously observed associations between metabolic rate or circulating glucocorticoids and life-history variation.

THs do not only regulate metabolisms in adults, but they are also transferred from mothers to developing offspring in all vertebrates (Ruuskanen and Hsu 2018). Such maternally-derived THs (hereafter “maternal THs”) play a critical role for early embryonic development (Moog et al. 2017; Darras 2019). In humans, the levels of maternal THs are associated with new-born body mass and children’s IQ (Medici et al. 2013; Korevaar et al. 2016). This suggests that variation in maternal THs can induce transgenerational phenotypic plasticity, which can constitute an important part of phenotypic variation in wild animals (Mousseau and Fox 1998; Moore et al. 2019). Nevertheless, the ecological and evolutionary significance of maternal THs in wild animals has been almost totally neglected (Ruuskanen and Hsu 2018). Recently, several experimental studies manipulating maternal THs reported variable effects of yolk TH on nestling traits across species (Ruuskanen et al. 2016a; Hsu et al. 2017, 2019a). These intriguing species differences suggest species might respond differently to elevated yolk TH levels and thus that maternal yolk THs might have co-evolved with life-history differences.

In this study, we explored the potential associations between maternal yolk TH levels and life-history strategies across 34 bird species. In birds, two maternally derived THs – triiodothyronine (T3) and thyroxine (T4) are transferred and stored in the egg yolk (Sechman and Bobek 1988; Prati et al. 1992). We asked whether the inter-specific variation of yolk THs would be associated with life-history traits across species, while accounting for environmental factors that may potentially contribute to the variation in yolk THs. The associations between maternal THs and life-history traits, if they exist, probably reflect the involvement of maternal THs in the evolution of life-history differences or vice versa.

Based on the known functions of THs, we considered three life-history continuums for which we hypothesized that maternal yolk THs may be associated with the related life-history traits. The first is the altricial-precocial developmental spectrum. At the two ends of this life-history continuum, precocial species and altricial species show distinctive ontogenetic patterns of visual and locomotory ability (Starck and Ricklefs 1998c), hypothalamic-pituitary-adrenal (HPA) axis (Wada 2008), hypothalamic-pituitary-thyroid (HPT) axis (McNabb 2006; de Groef et al. 2013) and thermoregulation (Price and Dzialowski 2018). In precocial species, the thyroid function already starts in the middle of incubation and the TH levels in thyroid gland and blood circulation both peak around hatching. By contrast, the thyroid function of altricial species starts days after hatching, and their circulating blood TH levels slowly increase towards the adult level (McNabb 2006; de Groef et al. 2013). Therefore, we hypothesized that such differences in the mode of development may be associated with yolk TH levels, either because higher maternal THs are required for a comparatively advanced development at hatching or because the developmental modes at least partly determine the transfer process of maternal THs, or both.

Second, we considered the life-history differences between migratory and resident species. THs have been known to regulate seasonal physiology and reproduction (Dardente et al. 2014; McNabb and Darras 2015; Nishiwaki-Ohkawa and Yoshimura 2016) and also moulting and feather regeneration (Vézina et al. 2009; Pérez et al. 2018). Several studies have also considered THs as part of the hormonal mechanism underlying migration (Pérez et al. 2016) and in general migratory species have higher BMR than resident species (Jetz et al. 2008). We thus hypothesized that maternal TH levels may be associated with a species’ migratory status.

Third, we considered the slow-fast pace-of-life (POL) continuum (Ricklefs and Wikelski 2002; Dammhahn et al. 2018). As introduced above, metabolic rates (Wikelski et al. 2003; Auer et al. 2018), glucocorticoids (Hau et al. 2010), and oxidative stress (Vágási et al. 2018) have been proposed to mediate the POL-related life-history variation. Interestingly, THs appear to have associations with all of them. It is well known that THs have regulatory function on metabolism (Mullur et al. 2014), and in wild birds, within-species studies have confirmed positive correlations between blood THs and BMR and RMR (e.g. Chastel et al. 2003; Elliott et al 2013; Welcker et al. 2013). THs also have cross-talks with the HPA axis (Cortés et al. 2014) and may induce higher oxidative stress due to the metabolic effects (Villanueva et al. 2013). Therefore, it seems possible that THs may also be a mediator of pace of life.

In addition to life-history differences, some environmental factors may also contribute to the inter-specific variation of maternal yolk THs and should be taken into account. First, we expected that the foraging environment, through its direct influence on food composition, may influence the inter-specific variation of maternal yolk THs. For example, food conditions before and during the egg-laying of captive wild-type rock pigeons (*Columba livia livia*) affected maternal yolk TH levels (Hsu et al. 2016). A possible explanation for this finding is the iodine content in the diet, as iodine is an essential ingredient for TH production (Bizhanova and Kopp 2009). In humans, even mild iodine deficiency during pregnancy has been shown to lead to deficiency of maternal T4 and hence detrimental effects on foetal development (Bath et al. 2013; Trumpff et al. 2013). In wild animals, environmental iodine availability is associated with antler weights of roe deer (*Capreolus capreolus*) via its influence on TH production (Lehoczki et al. 2011), and freshwater alligators (*Alligator mississippiensis*) have on average lower plasma T3 and T4 concentrations than their estuarine conspecifics (Boggs et al. 2011). Therefore, we hypothesized that iodine availability from the environment and diet (e.g. Eckhoff and Maage 1997) could contribute to part of the variation in maternal yolk THs. Moreover, captive environments differ in many aspects from wild environments, such as in food availability and safety from predators. Confinement and regular human disturbance might also pose as chronic stressors, which may influence THs (e.g. Angelier et al. 2016) and therefore maternal TH deposition. We therefore hypothesized that captivity may also influence maternal yolk THs.

To test these hypotheses, we collected information on life-history traits pertaining to each hypothesis from the literature. The environmental variables were assigned for each species based on the sampled population. We measured maternal THs in egg yolks, constructing the first interspecific maternal TH data set. By combining data of life-history traits and maternal THs, we were able to run phylogenetic mixed-effects models using a Bayesian framework to test our hypotheses.

## Materials and Methods

### Sample collection

We collected unincubated eggs from 34 species of birds across 17 families and 6 orders (n = 1-21 per species, median=7, Table S1). Nests of wild species were located by extensive nest searches in the known breeding habitats of each species or from nest-box populations during 2016-2017. We aimed at collecting eggs from species with varying body sizes and life-history traits and included both precocial and altricial species. All eggs were collected before clutch completion (i.e. when found, the eggs were cold and clutch was not completed) and one, randomly sampled egg per clutch was collected. Therefore, in most of the cases, the laying order of the sampled egg was unknown. Most of the wild species were sampled from Finnish populations near Turku (60°27’N, 22°16’E), Oulu (65°01’N, 25°28’E) and Kauhava (63°06’N, 23°04’E) regions, except thorn-tailed rayaditos (*Aphrastura spinicauda*, sampled mostly from Chiloé island, Chile, 42°40’S, 73°59’W) and collared flycatchers (*Ficedula albicollis*, from Burgsvik, Gotland, Sweden, 57°02’N, 18°16’E). The eggs of captive homing pigeons (*C. l. domesticus*) and domesticated chicken (*Gallus domesticus*) and eggs of semi-captive pheasants (*Phasianus colcichus*) and grey partridges (*Perdix perdix*) were sampled from local breeders in Finland. The eggs of red jungle fowl (*G. g. gallus*) and rock pigeons (*C. l. livia*) were sampled from the captive colonies at the University of Groningen, the Netherlands, and the eggs of captive zebra finches (*Taeniopygia guttata*) were from the University of Glasgow, UK. All collected eggs were first frozen and stored at −20°C until further analysis. For most of the species, the frozen eggs were shipped to the University of Turku, where we separated the yolk for hormone analysis (see *Yolk TH analysis*). Two exceptions are the collared flycatchers and the rock pigeons, for which the eggs had been dissected and the yolk had been homogenized and mixed in known amount of milli-Q (MQ) water before shipped to University of Turku and were already ready for extraction. Eggs of wild species were collected under licenses from the environmental authorities in each country: Finland, VARELY/665/2016, POPELY/61/2016, VARELY/63/2016, VARELY/412/2016, VARELY/6085/2016, and Finnish Wildlife Agency 84/2016; Chile, Agricultural Ministry 404/2017, Sweden, National Board for Laboratory Animals. Captive species were housed and eggs were collected following all national and international guidelines.

**Table 1.** Model, phylogeny and data sets defined in this study

| Model set | Phylogeny | Data set | Model structure (fixed factors) | Remark |
| --- | --- | --- | --- | --- |
| 1 | All 34 species | Full data set | ~developmental mode + migration + body mass + maximum lifespan + foraging environment + captivity |  |
| 2a | Excluding *Coturnix japonica* * | Excluding domesticated Japanese quails and domesticated chicken | ~developmental mode + migration + body mass + clutch size + clutches per year + captivity | Domesticated quails and chicken do not lay clear “clutches” of eggs. |
| 2b | Excluding *Coturnix japonica* | Excluding Japanese quails | ~developmental mode + migration + body mass + PC1 and PC2 of developmental period + captivity | Time to fledging data was missing for Japanese quails |
| 3 | Excluding *Gallus gallus* | Excluding red jungle fowl and domesticated chicken | ~developmental mode + migration + body mass + age at first reproduction + captivity | Age at first reproduction data was missing for both jungle fowl and chicken |
| 4 | 28 species with BMR data | Species with BMR data | ~developmental mode + migration + body mass + BMR + captivity |  |
| 5 | 29 species with growth rate data | Species with growth rate data | ~developmental mode + migration + body mass + growth rate + captivity |  |

\* In Model set 2a, both Japanese quails and domesticated chicken were removed from the “data set” but only *Coturnix japonica* was removed from the phylogeny because *Gallus gallus* was still needed in the phylogeny for the data from red jungle fowl.

### Yolk TH analysis

We extracted THs from the egg yolks and measured two THs (T3 and T4) at the University of Turku and Turku Centre of Biotechnology. Yolk was separated from albumen after a short period of thawing. A few eggs showed signs of very early development and all such eggs were discarded from the analysis. After dissection, yolks were weighed (∼0.01 g), and a half (for passerines) or a quarter (for other species) of the yolk was weighed for extraction and we added an equal amount of MQ to facilitate homogenizing.

We used the extraction protocol described previously (Ruuskanen et al. 2016b). In brief, we first added 2 ml methanol to the yolk-MQ mixture (ca. 300 mg) and homogenized the sample. A known amount of ^13^C_12_-T4 (Larodan) was added to each sample as an internal tracer, which allowed us to correct for extraction efficiency (i.e. recovery). Four ml of chloroform was then added and samples were centrifuged at +4°C 1900g for 15 min. The supernatant was then collected and the pellet was re-extracted in 2:1 chloroform-methanol mixture, back-extracted with 0.05% CaCl_2_ into aqueous phase, and re-extracted with chloroform:methanol:0.05% CaCl_2_ mixture (3:49:48). The aqueous phase was further purified on Bio-Rad AG 1-X2 resin columns, eluted with 70% acetic acid, and then vacuum-evaporated overnight. Samples from different species were spread over extraction batches and the extraction batch ID was included as a random intercept in the statistical models.

We used a validated nano-flow liquid chromatography-mass spectrometry (LC-MS) protocol to measure yolk T3 and T4 simultaneously (Ruuskanen et al. 2018). In brief, the dry TH extracts were re-suspended in 150 µl 0.1% NH_3_ and further diluted with 0.01% NH_3_. The dilution factor depended on the expected amount of T3 and T4 in each sample. Internal standards ^13^C_6_-T3 and ^13^C_6_-T4 were added to identify and also quantify T3 and T4 in the samples. A triple quadruple mass spectrometer (TSQ Vantage, Thermo Scientific, San Jose, CA) and nano-flow HPLC system Easy-nLC (Thermo Scientific) were used for the measurements. The on-column quantification limits were 10.6 amol and 17.9 amol for T4 and T3, respectively (Ruuskanen et al. 2018). The data was acquired automatically using Thermo Xcalibur software (Thermo Fisher Scientific) and subsequently analysed by Skyline (MacLean et al. 2010). TH concentrations were calculated using the peak area ratios of sample to internal standard, corrected for recovery, and expressed as pg/mg yolk. We further calculated the total amount of THs per yolk and expressed it as ng/yolk.

### Life-history traits and environmental factors

#### I. Altricial-Precocial continuum (i.e. developmental mode)

We specified each species as precocial or altricial, following Starck and Ricklefs (1998a). The four semi-precocial species, which all belong to the family Laridae (gulls and terns) and one semi-altricial species (the kestrel, *Falco tinnunculus*) were pooled with precocial or altricial species, respectively.

#### II. Migratory status

Each species was categorized as either migratory or resident, based on data compiled in Lehikoinen et al. (2003), McNab (2009) and Pap et al. (2015). When needed, the migratory status assignment was made based on the population from which we collected eggs.

#### III. Pace-of-life continuum

The pace-of-life continuum incorporates many related life-history traits, including a species’ developmental duration, fecundity, metabolic rate, growth rate, and the rate of maturation and senescence.

i. The data on **developmental duration** was collected mainly from the amniote life-history database compiled by Myhrvold et al. (2015) with supplementation from other literature (see *supplementary materials*). We collected the **incubation duration** and **the time to fledging** (from hatching) to represent the prenatal and postnatal part of the developmental period. We additionally calculated two more variables: **the total length of development** (incubation duration + time to fledging) and **the proportion of prenatal development** (incubation duration / total developmental length).
ii. The **growth rate** data originated from Starck and Ricklefs (1998b) and was supplemented with other literature (see *supplementary materials*). For comparability, the logistic growth rate constants were used to represent the growth rate of each species. Whenever multiple sources of the growth rates are available for a species, an average was used.
iii. **Species’ body mass, clutch size** and **number of clutches per year** were obtained from Myhrvold et al. (2015).
iv. **Basal metabolic rates** (BMR) were mainly obtained from McKechnie et al. (2006) and McNab (2009), supplemented by other literature (see *supplementary materials*).
v. **Maximum lifespan** and **ages at first reproduction** originated from the database AnAge (Tacutu et al. 2013), except for the thorn-tailed rayaditos (*Aphrastura spinicauda*), for which the data was obtained from Moreno et al. (2005) and Quirici et al. (2019).

#### IV. Environmental factors

For environmental factors, we created two variables to represent a species’ **foraging environment** and **captivity status.** Respectively, species were categorized as marine-bound (foraging on marine food resources) or terrestrial (foraging on non-marine bound resources) based on the breeding territory and diet, and wild or captive. Semi-captive species (the pheasant and the grey partridge) were categorized as captive.

All life-history and environmental data we compiled for this study, the sources we consulted, and the final values used in all our models, can be found in the supplementary materials.

### Phylogenetic mixed models

#### Model specifications

Following Hadfield and Nakagawa (2010), we ran phylogenetic mixed models with the R package *MCMCglmm* (Hadfield 2010). T3 and T4 concentrations (pg/mg yolk) as well as total contents (ng/yolk) were ln-transformed and standardized and used as dependent variables in separated univariate models. Species body mass and other life-history and environmental variables (see *Life-history traits and environmental factors*) were treated as fixed factors and covariates. In order to facilitate model fitting, body mass was also ln-transformed and standardized. Before fitting models, all two-level categorical variables were first dummy-coded as −0.5 and 0.5 (Table S2), to facilitate model mixing/convergence. The life-history traits that are known to be associated with body mass (BMR, growth rate, maximum lifespan, age at first reproduction) were first corrected for body mass and phylogenetic relatedness among species using the R package *phytools* (Revell 2009, 2012). BMR data, in particular, was first converted to mass-specific BMR (KJ/h/g) and ln-transformed and corrected for body mass and phylogeny.

**Table 2.** Phylogenetic signals (i.e. phylogenetic heritability) for maternal yolk T3 and T4.

| Hormone | Parameter | Posterior mean [95% CI] |
| --- | --- | --- |
| T3 | concentration | 0.84 [0.68, 0.95] |
|  | total content | 0.84 [0.71, 0.95] |
| T4 | concentration | 0.60 [0.25, 0.87] |
|  | total content | 0.75 [0.54, 0.91] |

The four variables of developmental duration were strongly correlated with one another. Therefore, we conducted a phylogeny-corrected principal component analysis (PCA) using R package *phytools* (Revell 2009, 2012). The first two PCs explained 99.74% of the variance (Table S3). The loadings suggested that PC1 is highly positively correlated with the time to fledging and the total length of developmental period, and negatively correlated with the proportion of prenatal development, while PC2 is negatively correlated with the duration of prenatal development (Table S3). In subsequent phylogenetic mixed models, we therefore used the scores of PC1 and PC2 for each species to represent the developmental duration.

Because the data of life-history traits were not available for all species, we defined several sets of models depending on data availability, listed in Table 1. Each model and data set was thus pruned for the specific set of species where data was available. Each model set consists of separate models to analyse the concentrations and total contents of T3 and T4, respectively. This is because in oviparous species, the total contents of maternal hormones in the yolk is fixed after yolk formation. Therefore, the total content of hormones represents an initial maximum hormone availability and may represent different biological significance from the concentration.

In all models, the phylogenetic relatedness among all species included was treated as random effects, following Hadfield and Nakagawa (2010). Additionally, species name was also included as a random factor to control for within-species non-independence among the multiple eggs from the same species. For this random factor, homing pigeons and domesticated chickens were coded separately from their wild-type ancestor rock pigeons and red jungle fowl, despite that they were not distinguished in the phylogeny (i.e. coded with the same scientific names, *Columba livia* and *Gallus gallus*). The batch of hormone extraction was also included as an additional random factor. Response variables were all assumed to follow Gaussian distributions. All models were run for 750000 iterations with a 350 thinning interval and burn-in of 50000, aiming for an approximately 2000 effective sample size for all parameters. Following Houslay and Wilson (2017), we used an uninformative parameter-expanded prior following a Cauchy distribution (V=1, nu=1, alpha.mu=0, alpha.V=25^2) for all random factors. Tests using an uninformative inverse-Wishart prior or improper flat prior gave qualitatively same results, suggesting the results are robust against the choice of priors.

#### Phylogenetic trees and phylogeny uncertainty

Each phylogeny set (Table 1) was correspondingly pruned and obtained from BirdTree.org (Jetz et al. 2012) based on two phylogenetic backbones (Ericson et al. 2006; Hackett et al. 2008). For each backbone, 100 possible trees were generated and downloaded. All models were tested repeatedly across the 100 possible trees in each backbone and the posterior means of parameter estimates and their 95% credible intervals (CIs) were stored. The average posterior mean and the average boundaries of the 95% CIs were calculated to account for phylogenetic uncertainty (Rubolini et al. 2015) and reported. The results are highly similar across all possible trees (see Fig. S1 for examples) and between the two backbones. Therefore we only reported the results based on the Hackett backbone.

#### Model diagnostics

As models across all possible phylogenetic trees produced highly similar results, we examined model performance by re-fitting all models with a consensus tree. The consensus tree was derived using the R package *phytools* (Revell 2012) for each phylogeny set. Model diagnostics were conducted by visual inspection on the trace plots for proper mixing and on autocorrelations. Model convergence was also checked using Gelman diagnostics provided in the R package *coda* (Plummer et al. 2006). For all models, visual inspection did not raise any red flags for poor mixing or substantial autocorrelation and all models passed the Gelman diagnostics with all potential scale reduction factors < 1.05.

#### Phylogenetic signals

Phylogenetic signal, a.k.a. phylogenetic heritability (*H^2^*), was calculated as

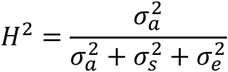

where 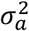 represents the variance of the phylogeny; 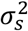 represents the variance accounted by individual species; 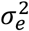 represents the residual variance. The *H^2^* calculated this way is equivalent to the Pagel’s λ (Hadfield and Nakagawa 2010).

Because phylogenetic signal is in fact the proportion of the phylogenetic variance over the total phenotypic variance, the fixed factors included in the model will influence the value of the phylogenetic signal by changing the estimates of all variance components (Nakagawa and Schielzeth 2010). Therefore we calculated the phylogenetic signals for yolk T4 and T3 based on the phylogenetic mixed model that only included the life-history traits whose 95% CIs did not encompass 0 (i.e. having credible associations). These are the developmental mode, migratory status, captivity, and body mass (see *Results*). In these models, all 34 species and the full phylogeny set was included and we used a consensus tree from the Hackett backbone, derived by using the *phytools* package (Revell 2012).

## Results

### Inter-specific variation in yolk THs and phylogenetic signals

We observed substantial inter-specific variation in both T3 and T4 concentrations (Fig. 1, Table S1). Great tits (*Parus major*) and redshanks (*Tringa totanus*) had the lowest and highest concentrations of yolk T3 and T4 among the 34 species, respectively. Between the two species, the difference in average yolk THs reached 100-fold for T3 (great tit, mean ± SD = 0.112 ± 0.032 pg/mg yolk, n = 11; redshank, mean ± SD = 11.242 ± 5.067 pg/mg yolk, n = 4), and 18-fold for T4 (great tit, mean ± SD = 0.989 ± 0.292 pg/mg yolk, n = 12; redshank, mean ± SD = 18.101 ± 4.125 pg/mg yolk, n = 4).

**Figure 1.**
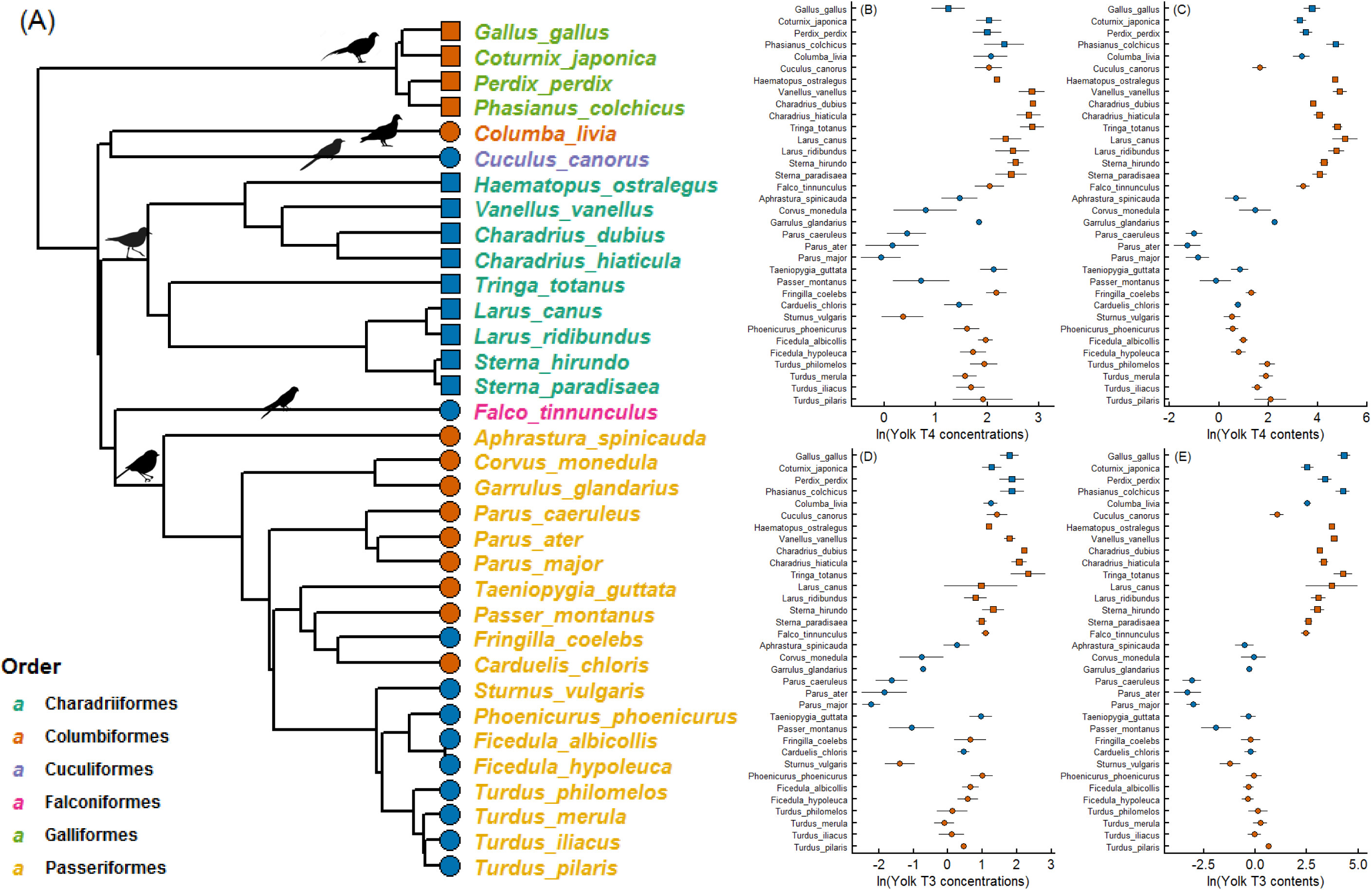
Phylogenetic tree, yolk TH concentrations and total contents of the avian species included in this study. The phylogenetic tree (A) is one possible tree derived from the Hackett backbone (see text). Different colours in species names represent different orders they belong to. Yolk TH concentrations (B, D) and total contents (C, E) exhibited substantial inter-specific variation (mean ± SD, see Table S1 for exact values and sample sizes). Red symbols represent resident species, while blue symbols migratory species. Circles represent altricial species, and squares precocial species.

The moderate to strong phylogenetic signals indicated that the phylogenetic relatedness among species accounted for 60%-85% of the inter-specific variation in yolk THs (Table 2). On average, T3 has stronger phylogenetic signals than T4, although their 95% CIs overlapped.

### Associations between yolk THs and life-history variables

The estimated posterior means and 95% CIs from all models are presented in Fig. 2. Precocial species deposited larger total amounts of THs in the egg yolks than altricial species (Fig. 3), but not higher TH concentrations. The model set 1 supported this difference with positive posterior means and 95% CIs, when controlling for species body mass and other covariates, especially for T3 contents (Fig. 2D). Also, migratory species generally deposited higher concentrations and also larger amounts of both THs in the egg yolks than resident species (Fig. 4), which were supported by the model set 1 (Fig. 2). Species body mass also strongly correlated with yolk TH content, but not with yolk TH concentration (Fig. 2). Along the slow-fast pace-of-life continuum, however, no traits were found to be credibly associated with maternal yolk TH concentrations or contents (Fig. 2)

**Figure 2.**
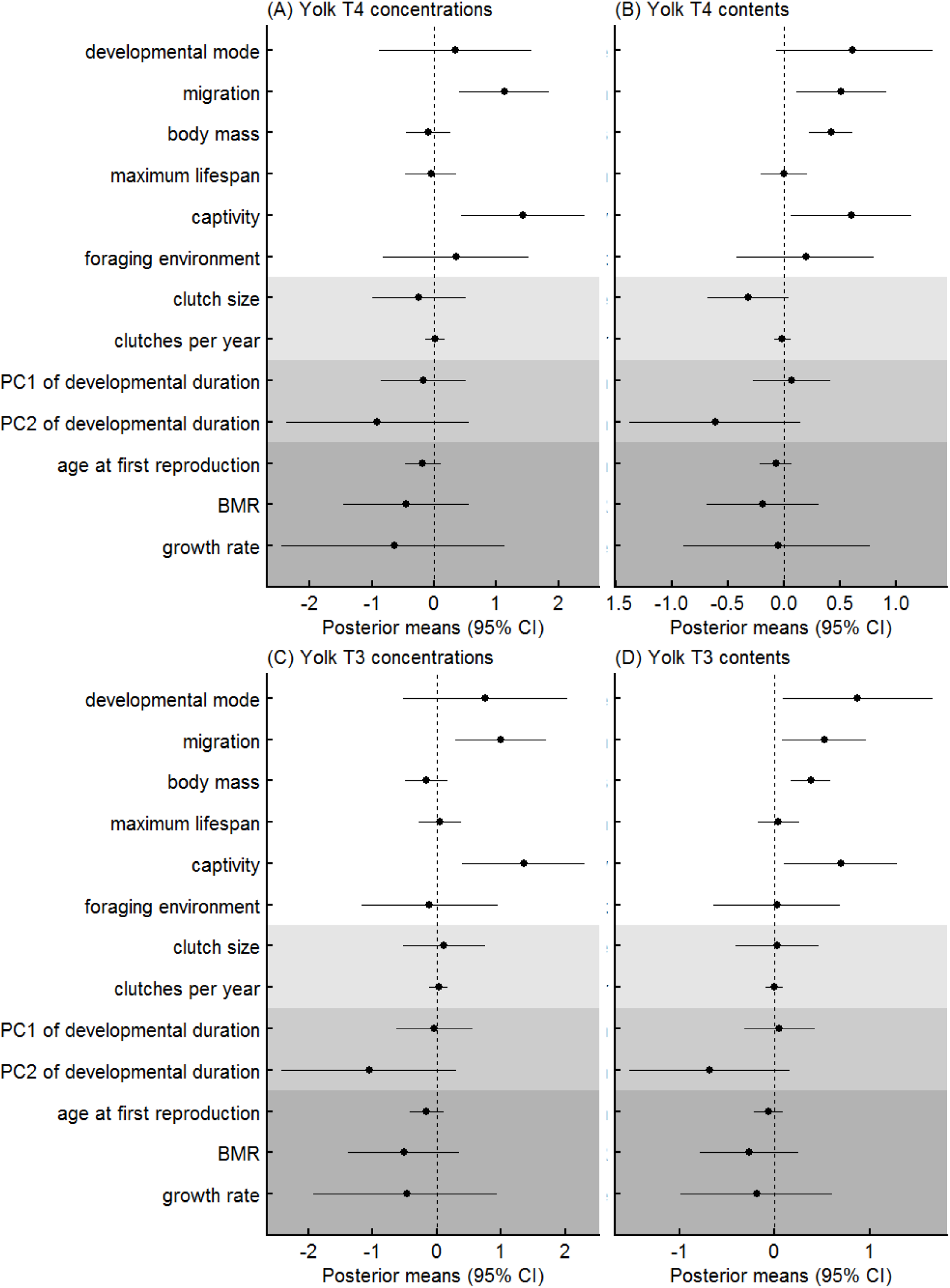
Posterior means (± 95% credible intervals) between yolk THs and all life-history variables tested in this study. White area presents the estimates from model set 1 (see text and Table 1). The shaded areas present the variables tested in model set 2 (the light and medium grey areas) and model set 3-5 (the darkest grey area). In models 2-5, the estimates of the variables that had been tested in model set 1 are not redundantly presented. For developmental mode, the estimate represents the difference of yolk THs in precocial species from altricial species. For migration, the estimate represents the difference in migratory species from resident species. For captivity, the estimate represents the difference in captive species from wild species. For foraging environment, the estimate represents the difference in marine-bound species from terrestrial species.

**Figure 3.**
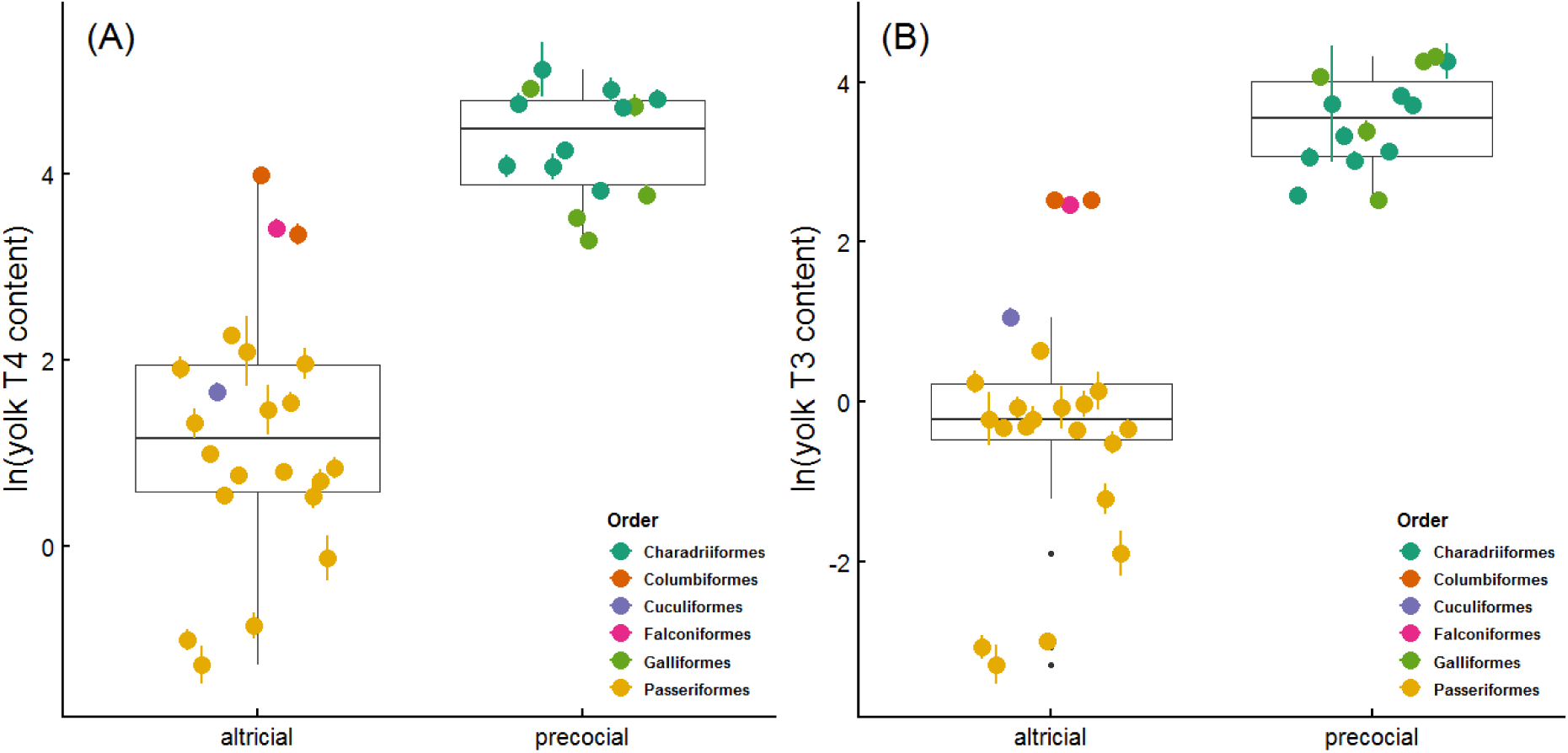
Boxplots and species-specific averages of yolk T4 (A) and T3 (B) contents (ng/yolk, ln-transformed raw data) according to developmental mode. Boxplots represent the median (the middle line) and the first and the third quartiles (the box), and the whiskers extend to 1.5 times of the interquartile range. Colored dots represent species-specific means (±SE).

**Figure 4.**
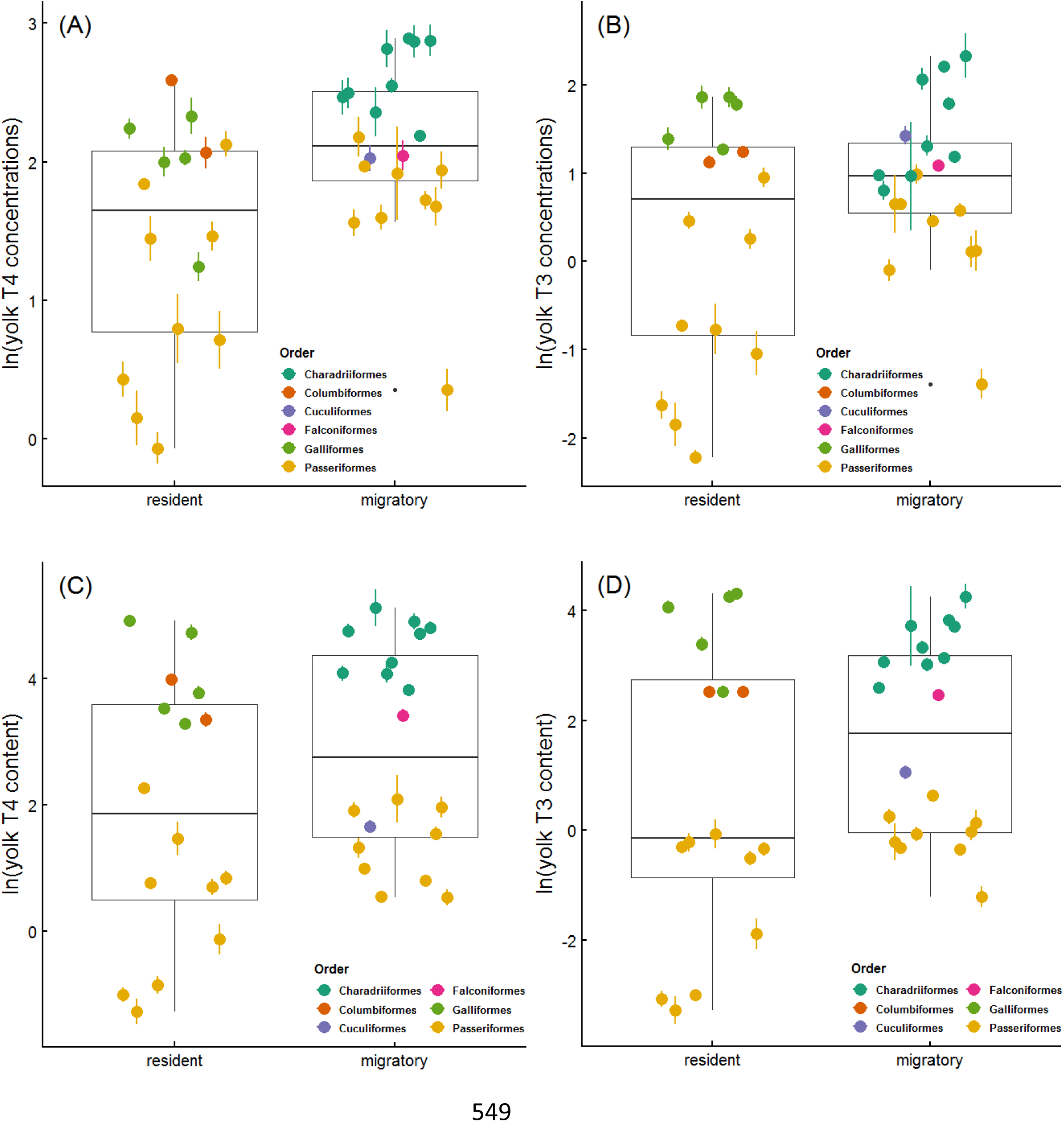
Boxplots and species-specific averages of yolk TH concentrations (A, B, pg/mg, ln-transformed raw data) and total contents (C, D, ng/yolk, ln-transformed raw data) between migratory and resident species. Boxplots represent the median (the middle line) and the first and the third quartiles (the box), and the whiskers extend to 1.5 times of the interquartile range. Colored dots represent species-specific means (±SE).

### Associations between yolk THs and environmental variables

Although the raw data suggested that marine-bound species deposited larger amounts of yolk THs than terrestrial species (Fig. S2), this was not supported by the phylogenetic mixed models (Fig. 2). Between captive and wild species, credible differences in both yolk T3 and T4, and in both concentrations and total contents (Fig. 2) were found. This suggested that captive species deposited higher concentrations and larger amounts of yolk THs than wild species (Fig. 5).

**Figure 5.**
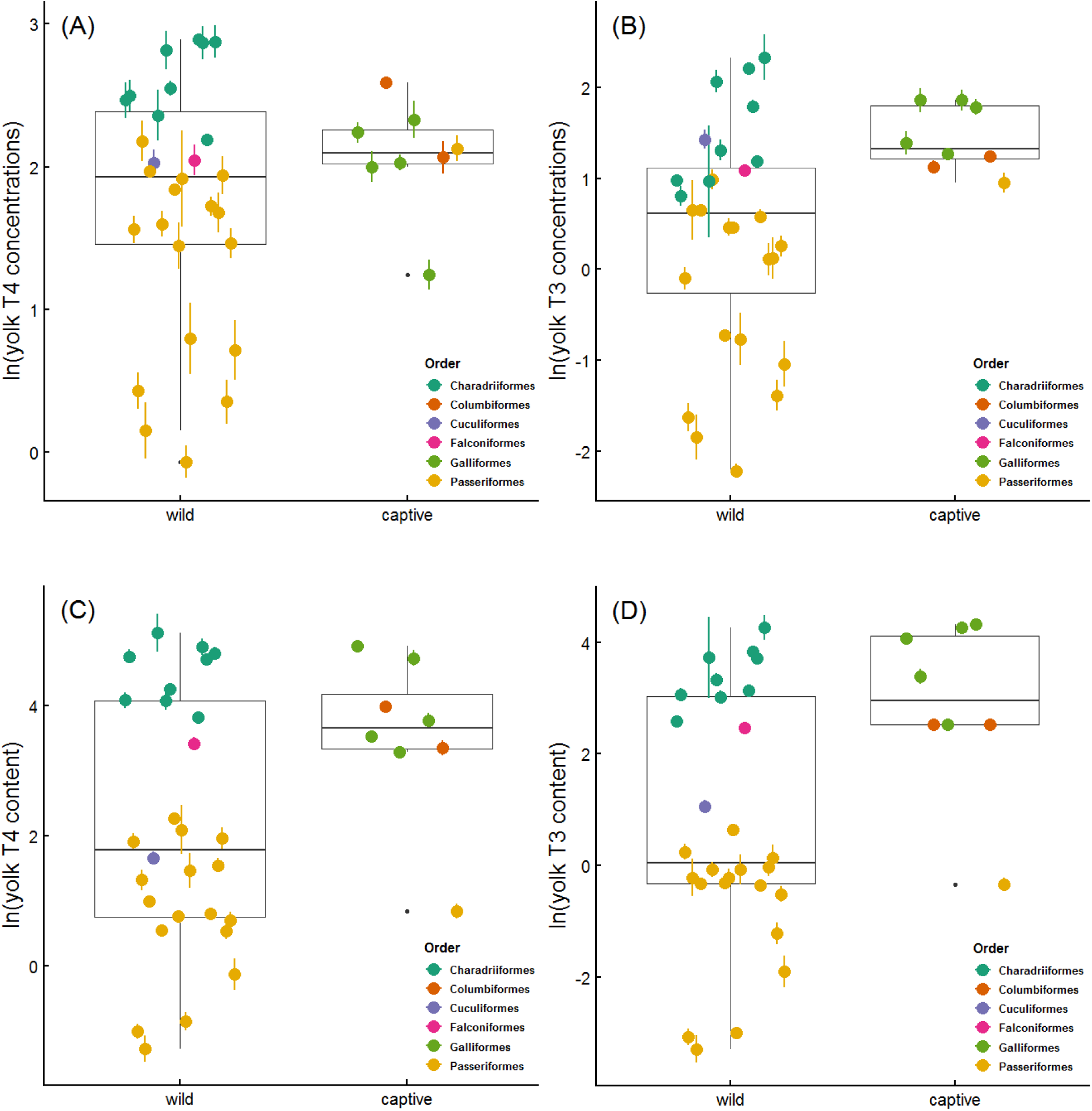
Boxplots of yolk TH concentrations (A, B, pg/mg, ln-transformed raw data) and total contents (ng/yolk, ln-transformed raw data) in between captive and wild species. Boxplots represent the median (the middle line) and the first and the third quartiles (the box), and the whiskers extend to 1.5 times of the interquartile range. Colored dots represent species-specific means (±SE).

## Discussion

### Maternal THs and life-history variation

Among the three life-history continuums, our phylogenetic mixed models estimated credible differences in maternal yolk THs between the migratory and resident life histories, and between precocial and altricial developmental modes. However, contrary to our expectation, none of the pace-of-life (POL) related traits were found credibly associated with maternal yolk THs, after controlling for body mass. The moderate to strong phylogenetic signals also indicate that a substantial proportion of variation in maternal THs is accounted for by the phylogeny itself. This suggests a substantial extent of phylogenetic inertia, which may present some evolutionary constraints in maternal TH deposition. Despite the constraint, the credible differences of maternal THs between precocial and altricial species, and between migratory and resident species still suggest that maternal THs are associated with the evolution of these two life-history continuums or vice versa.

### THs and migration

Migration is a critical and demanding life-history stage. Many physiological changes occur both before and after migration, such as fat deposition, gonadal regression and recrudescence (Hahn et al. 2015). The temporal overlap with other seasonal activities, such as breeding and moult, also tends to be minimized over evolutionary history in migratory species (Hahn et al. 2015). By contrast, resident species usually have a relatively longer time window to coordinate the physiological transition before and after reproduction compared with migratory species. Therefore, migratory species may require a higher sensitivity or accuracy to fine-tune the transition. THs play a central role in the molecular mechanisms of photoperiodic response. The photoperiodic change influences the expression levels of the thyroid-stimulating hormones (TSH) in the hypothalamus, which subsequently induces reciprocal switching of two deiodinases (DIO2, converting T4 to active T3 and DIO3, converting T4 and T3 to inactive THs) that activate and inactivate T3, respectively (Nakao et al. 2008; Dardente et al. 2014). This photoperiodic DIO2/DIO3 expression pattern further regulates the seasonal maturation of gonads. Interestingly, the photoperiodic DIO2/DIO3 gene expression pattern of great tits was only evident in a Swedish population but not in a southern German population (Perfito et al. 2012), in which the duration of the breeding seasons are likely to differ. Our result of higher yolk TH levels in migratory than resident species, not only in terms of higher concentrations but also larger amounts, is therefore in line with the notion that the differences in maternal THs could have some organizational effects on TH responsiveness later in life. There may still be species differences in the seasonal regulation of DIO2/DIO3 expression, as a similar pattern was not found in European starlings (*Sturnus vulgaris*, Bentley et al. 2013). Comparison of TH-related gene expression between conspecific populations that possess different migratory propensity, potentially in combination with TH manipulation, may help to identify the elements responsible for such differences.

#### Does precocial development require more maternal THs?

We found that precocial species deposited higher amounts of THs (accounted for body mass), particularly T3, in the egg yolks than altricial species, which fits their distinct developmental trajectories (Starck and Ricklefs 1998c). Compared with altricial species, precocial species temporally shift their physiological maturity to an earlier life stage in terms of visual and locomotory ability (Starck and Ricklefs 1998c), HPA and HPT axis (McNabb 2006; Wada 2008; de Groef et al. 2013) and thermoregulation (Price and Dzialowski 2018). Our result may suggest the requirement of a larger amount of maternal THs for such an advanced ontogenetic trajectory in precocial birds.

An alternative way to look at the contrasting ontogenetic differences between precocial and altricial development may be the placement of hatching on the ontogenetic timeline. Regardless of the developmental modes, the event of hatching requires a lot of physiological changes to prepare the chick for the “outside” world and therefore marks a major life-stage transition. As the altricial developmental mode is assumed to have evolved later than the precocial mode (Starck and Ricklefs 1998c; de Groef et al. 2013), one way to look at the shift is that the event of hatching is moved earlier along the ontogenetic timeline in the altricial development so that hatching occurs when the young is still poorly developed. Whether maternal THs mediate such a shift during the evolution from precocial to altricial development requires further studies, but according to our data such a change may coincide with a reduction in TH deposition over the evolutionary course. We therefore could expect intermediate levels of maternal THs in semi-precocial and semi-altricial species. Interestingly, the only semi-altricial species we measured, the Eurasian kestrel, indeed has higher maternal TH content than most other altricial species, but lower than most semi-precocial and precocial species (Fig. 3).

#### No size-independent associations between maternal THs and pace of life (POL)

Given the known function of THs on metabolism and interactions with glucocorticoids, it is surprising that we did not detect any associations between maternal THs and all tested POL-related traits (Fig. 2). Although yolk TH contents show strong correlations with species body mass, this is because large species produce large eggs and large yolks, hence higher total amounts of yolk THs. After controlling for body mass and phylogenetic relatedness, only clutch size showed a potential trend with T4 contents. The model estimated a negative posterior mean and the 95% CI was slightly overlapped with 0 (Fig. 2B), implying that species with larger clutch sizes tended to deposit less T4 in the egg yolks. Such a trend is probably driven by the difference between passerines and shorebirds (Fig. S3). Confirming such a relationship perhaps would require a larger sampling scheme that includes more species.

The lack of associations between maternal yolk THs and POL-related traits does not support our hypothesis that THs were mediators of POL. This result therefore implies that either (i) the effects of maternal THs may not be translated directly to blood THs or (ii) blood THs are not associated with POL-related traits. The former could be due to limited organizational effects of maternal THs on the HPT axis or a more complex manifestation, such as sex-specific effects. To our knowledge, there are only few experimental studies that have examined how maternal THs influenced blood THs and metabolic rates during the nestling phase. In the great tit, yolk TH elevation did not influence nestling RMR before fledging (Ruuskanen et al. 2016a). In the rock pigeon, yolk TH elevation raised the RMR in female nestlings but reduced it in males on hatching day, a pattern that was consistent with the effects on plasma T4 concentrations (but not T3) at day 14 after hatching (in the middle of the nestling phase, Hsu et al. 2017). Although elevated yolk THs showed consistent effects on nestling RMR and plasma T4, the sex-specific pattern remains unexplained. Nevertheless, in domesticated chicken embryos the injection of THs into the yolk influenced the expression of two TH-transporters in the brain (van Herck et al. 2012). This suggests that differential exposure to maternal THs may influence brain TH availability. Such changes might have the potential to alter the sensitivity and responsiveness of the HPT axis in the long term. Therefore, the notion that maternal THs may organizationally shape the HPT axis cannot yet be ruled out. Another possibility to explain the lack of associations between maternal THs and POL-related traits is that blood THs are not associated with POL-related traits. This would be quite surprising considering the close associations between THs with metabolic rates, glucocorticoids, and oxidative stress. As POL-related traits are usually strongly associated with body mass, perhaps a much larger number of species is required to detect such potential associations after accounting for species body mass.

#### Maternal THs and life-history variation: cause or consequence?

Above, we have discussed the potential role of maternal THs as mediators of life-history variation. However, it is possible that differences in life-histories, particularly in migratory behaviour and developmental mode, may result in physiological differences that cause differential maternal TH deposition during egg formation across species. For example, migratory birds generally have higher BMR than residents (Jetz et al. 2008). Given the close relationship between BMR and THs, migratory species probably also have higher circulating blood TH levels. One study in two closely-related skylark species indeed found that the migratory Eurasian skylarks (*Alauda arvensis*) had higher pre-breeding blood T3 levels than the resident Asian short-toed larks (*Calandrella cheleensis*, Zhao et al. 2017). A meta-analysis on the available blood TH data from the literature should be promising to test the associations between life-history variation and TH physiology but to our knowledge, such a meta-analysis has not yet been done. If in general migratory species keep comparatively higher circulating TH levels than resident species, the higher maternal TH deposition could be simply mirroring the circulating TH levels in the system of migratory birds.

Even if maternal yolk THs are a physiological consequence induced by different life histories, differential yolk TH deposition may still reinforce the evolution of the life-history variation. Due to the critical role in embryonic development, maternal THs likely have short-term and long-term effects on offspring phenotype, such as nestling development, growth, and physiology (Ruuskanen et al. 2016a; Hsu et al. 2017, 2019a). If higher maternal THs would lend beneficial effects for a certain life-history strategy, and physiologically such a life history results in a higher maternal TH transfer, maternal THs may speed up the evolution of associated life-history traits (McGlothin and Galloway 2014). At the moment, this is a bold speculation. However, considering that yolk T3 deposition has a moderate heritability (h^2^ = 0.25, Ruuskanen et al. 2016c) and clear between-individual difference (Hsu et al. 2019b), there might be evolutionary potential for this to happen. Future studies may examine the temporal and quantitative differences in the expression patterns of TH-related genes in embryonic brains between migratory and resident species, and between precocial and altricial species, to know if there has been fundamental differences during the early developmental stage. Another angle would be examining whether birds hatching from eggs with elevated yolk THs may deposit higher yolk THs than control birds to test if maternal TH can induce any non-genetic inheritance.

### Maternal THs and environment

In terms of environmental factors, our models suggested differences in maternal yolk THs between captive and wild species, but not between terrestrial and marine-bound species.

#### Captive life has favoured higher maternal TH deposition

Life in captivity is characterised by space confinement, (usually) unlimited food availability and a predator-free environment (Mason 2010). The contrast to life in the wild can influence phenotypic expression (i.e. phenotypic plasticity) and even cause selection for certain traits (Driscoll et al. 2009). For example, captive zebra finches, domesticated or not, started incubating earlier and therefore induced a stronger degree of hatching asynchrony than wild zebra finches (Gilby et al. 2013). In our study, all the captive species have acclimated to captive lives for generations. Therefore the result that captive species deposit higher levels of THs in the egg yolks likely reflects some selection under captivity. The actual selective force is elusive at the moment, but probably related to traits like faster acclimation/habituation for anthropogenic diet or perhaps dampened stress response to confined space and constant human disturbance, which should be favoured in captivity.

#### Lack of evidence for the potential link with proximity to the sea

The larger yolk TH contents in marine-bound species compared to terrestrial species, apparent from the raw data, were not supported by the phylogenetic mixed models. Our data likely lacked the power to statistically partition the effects of environment and phylogeny because all sampled marine-bound species were from the order Charadriiformes. Future studies should therefore broaden the sampling scheme to overcome this problem. An alternative to test the relationship between foraging environment and maternal THs is to focus on a smaller clade, such as a single order or a single species inhabiting both inland and marine-bound populations. In the present study, the eggs of the little ringed plover (*Charadrius dubius*) were sampled from an inland population. The species, interestingly, had the lowest T4 content among all Charadriiform species, which implies possible influence from the environment. Experimentally manipulating the iodine content of diet in a captive model system or supplementing iodine-rich food to a wild population would be also an effective approach in this respect.

### Conclusion

Our phylogenetic mixed models suggested that migratory species deposited higher levels of both THs in the egg yolks than resident species, and precocial species deposited higher total content of T3 in the egg yolks than altricial species. In addition, captive species also consistently deposited higher levels of yolk THs, suggesting some selection process under captivity. These identified patterns suggest that maternal THs may be involved in the process of life-history evolution and can respond to on-going selective forces. Further effort should be invested to uncover the underlying physiological mechanisms.

## Acknowledgement

We thank Jon Brommer for his kind and helpful statistical advice in the beginning stage of data analysis. We thank Jorma Nurmi for excellent nest-finding skills and effort in the field. Mass spectrometry analyses were performed at the Turku Proteomics Facility, University of Turku and Åbo Akademi University. The facility is supported by Biocenter Finland. This study was funded by a grant from the Academy of Finland to SR (grant no. 286278).

