## supplementary materials for "Maternally-transferred thyroid hormones and life-history variation in birds"

**Table S1.** Mean ± SD of yolk T3 and T4 concentrations and total contents of all 34 species in this study.

| Species | N of T3 | Yolk T3 concentration (pg/mg yolk) | Yolk T3 content (ng/yolk) | N of T4 | Yolk T4 concentration (pg/mg yolk) | Yolk T4 content (ng/yolk) |
|---|---|---|---|---|---|---|
| Japanese quail, *Coturnix japonica* | 21 | 3.690±1.009 | 13.035±3.787 | 21 | 7.802±1.661 | 27.514±6.846 |
| Red jungle fowl, *Gallus gallus gallus* | 10 | 6.163±1.852 | 78.122±25.792 | 10 | 3.633±1.215 | 45.991±16.501 |
| Domesticated chicken, *Gallus gallus domesticus* | 5 | 4.151±1.315 | 59.809±16.053 | 5 | 9.488±1.621 | 137.149±16.649 |
| Grey partridge, *Perdix perdix* | 7 | 6.803±2.765 | 31.229±12.491 | 7 | 7.633±2.159 | 34.930±9.072 |
| Ringed-necked pheasant, *Phasianus colchicus* | 9 | 6.743±2.010 | 74.055±22.609 | 9 | 11.060±5.022 | 120.725±52.997 |
| Common cuckoo, *Cuculus canorus* | 8 | 4.322±1.065 | 2.998±0.788 | 8 | 7.821±2.112 | 5.397±1.462 |
| Rock pigeon, *Columba livia livia* | 9 | 3.532±0.785 | 12.685±2.351 | 9 | 8.303±2.982 | 29.978±10.289 |
| Homing pigeon, *Columba livia domesticus* | 4 | 3.103±0.463 | 12.575±2.084 | 4 | 13.339±0.687 | 53.981±3.341 |
| Eurasian oystercatcher, *Haematopus ostralegus* | 1 | 3.285 | 40.782 | 1 | 8.946 | 111.068 |
| Northern Lapwing, *Vanellus vanellus* | 5 | 6.051±0.942 | 46.195±6.027 | 5 | 18.050±4.497 | 139.166±39.438 |
| Little ringed plover, *Charadrius dubius* | 1 | 9.098 | 22.972 | 1 | 17.984 | 45.409 |
| Common ringed plover, *Charadrius hiaticula* | 3 | 8.021±1.630 | 28.325±5.727 | 3 | 16.965±3.871 | 59.980±14.134 |
| Redshank, *Tringa tetanus* | 4 | 11.242±5.067 | 76.245±32.722 | 4 | 18.101±4.125 | 123.911±25.503 |
| Common tern, *Sterna hirundo* | 8 | 3.883±1.250 | 21.563±7.313 | 8 | 12.920±1.844 | 71.466±11.647 |
| Arctic tern, *Sterna paradisaea* | 6 | 2.681±0.429 | 13.602±2.926 | 6 | 12.229±3.479 | 61.544±17.678 |
| Common gull, *Larus canus* | 3 | 3.723±3.521 | 67.738±75.481 | 3 | 10.929±3.444 | 182.853±99.314 |
| Black-headed gull, *Larus ribidundus** | 9 | 2.363±0.883 | 22.773±9.292 | 9 | 12.789±4.801 | 121.907±43.035 |
| Kestrel, *Falco tinnunculus* | 7 | 3.000±0.356 | 12.001±2.698 | 7 | 7.992±2.238 | 31.472±9.262 |
| Thorn-tailed rayadito, *Aphrastura spinicauda* | 11 | 1.375±0.503 | 0.656±0.278 | 11 | 4.564±1.647 | 2.192±0.996 |
| Jackdaw, *Corvus monedula* | 5 | 0.552±0.390 | 1.079±0.703 | 6 | 2.529±1.248 | 4.982±2.464 |
| Eurasian jay, *Garrulus glandarius* | 1 | 0.485 | 0.739 | 1 | 6.287 | 9.568 |
| Blue tit, *Parus caeruleus** | 9 | 0.217±0.104 | 0.051±0.024 | 9 | 1.647±0.659 | 0.385±0.134 |
| Great tit, *Parus major* | 11 | 0.112±0.032 | 0.053±0.020 | 12 | 0.989±0.292 | 0.458±0.162 |
| Coal tit, *Parus ater** | 7 | 0.186±0.106 | 0.045±0.027 | 7 | 1.301±0.619 | 0.312±0.157 |

| | | | | | | |
|---|---|---|---|---|---|---|
| Zebra finch, *Taeniopygia guttata* | 9 | 2.729±1.084 | 0.767±0.372 | 9 | 8.681±2.733 | 2.462±1.028 |
| Tree sparrow, *Passer montanus* | 7 | 0.415±0.232 | 0.187±0.127 | 7 | 2.294±1.046 | 1.015±0.510 |
| Chaffinch, *Fringilla coelebs* | 2 | 2.016±0.896 | 0.853±0.391 | 2 | 8.927±1.798 | 3.768±0.815 |
| Greenfinch, *Carduelis chloris** | 3 | 1.603±0.285 | 0.826±0.244 | 3 | 4.366±1.276 | 2.160±0.273 |
| European starling, *Sturnus vulgaris* | 7 | 0.273±0.131 | 0.334±0.179 | 7 | 1.537±0.695 | 1.794±0.670 |
| Common redstart, *Phoenicurus phoenicurus* | 8 | 2.805±0.850 | 0.998±0.367 | 8 | 5.073±1.151 | 1.774±0.424 |
| Collared flycatcher, *Ficedula albicollis* | 15 | 1.978±0.489 | 0.753±0.201 | 15 | 7.209±0.998 | 2.737±0.418 |
| Pied flycatcher, *Ficedula hypoleuca* | 15 | 1.862±0.570 | 0.740±0.238 | 15 | 5.763±1.421 | 2.307±0.654 |
| Song thrush, *Turdus philomelos* | 4 | 1.222±0.586 | 1.260±0.678 | 4 | 7.157±2.045 | 7.389±2.607 |
| Blackbird, *Turdus merula* | 6 | 0.939±0.307 | 1.353±0.525 | 6 | 4.881±1.233 | 7.013±2.147 |
| Fieldfare, *Turdus pilaris* | 3 | 1.587±0.111 | 1.904±0.257 | 3 | 7.521±3.852 | 9.200±5.259 |
| Redwing, *Turdus iliacus* | 4 | 1.168±0.376 | 1.009±0.276 | 4 | 5.509±1.543 | 4.745±0.996 |

* In order to match the phylogenetic trees from BirdTree.org, “old” scientific names were still used in this study. Their updated scientific names according to The Clements checklist of birds of the world v2018 (Clements et al. 2018) are: Chroicocephalus *ridibundus*, *Cyanistes caeruleus, Periparus ater, Chloris chloris*

**Table S2.** Dummy code for the two-level categorical variables included in the phylogenetic mixed models.

| | -0.5 | 0.5 |
|---|---|---|
| Mode of development | altricial | precocial |
| Migratory status | resident | migratory |
| Foraging environment | terrestrial | marine-bound |
| Captivity | wild | captive |

**Table S3.** Proportion of variance explained and loadings of the phylogeny-corrected PCA among the four variables of developmental duration.

| | **PC1** | **PC2** |
|---|---|---|
| Proportion of variance | 83.26 | 16.48 |
| Cumulative proportion | 83.26 | 99.74 |
| **Loadings** | | |
| Incubation duration | 0.264 | -0.965 |
| Time to fledging | 0.998 | 0.055 |
| Total developmental length | 0.942 | -0.329 |
| Prenatal proportion | -0.869 | -0.489 |

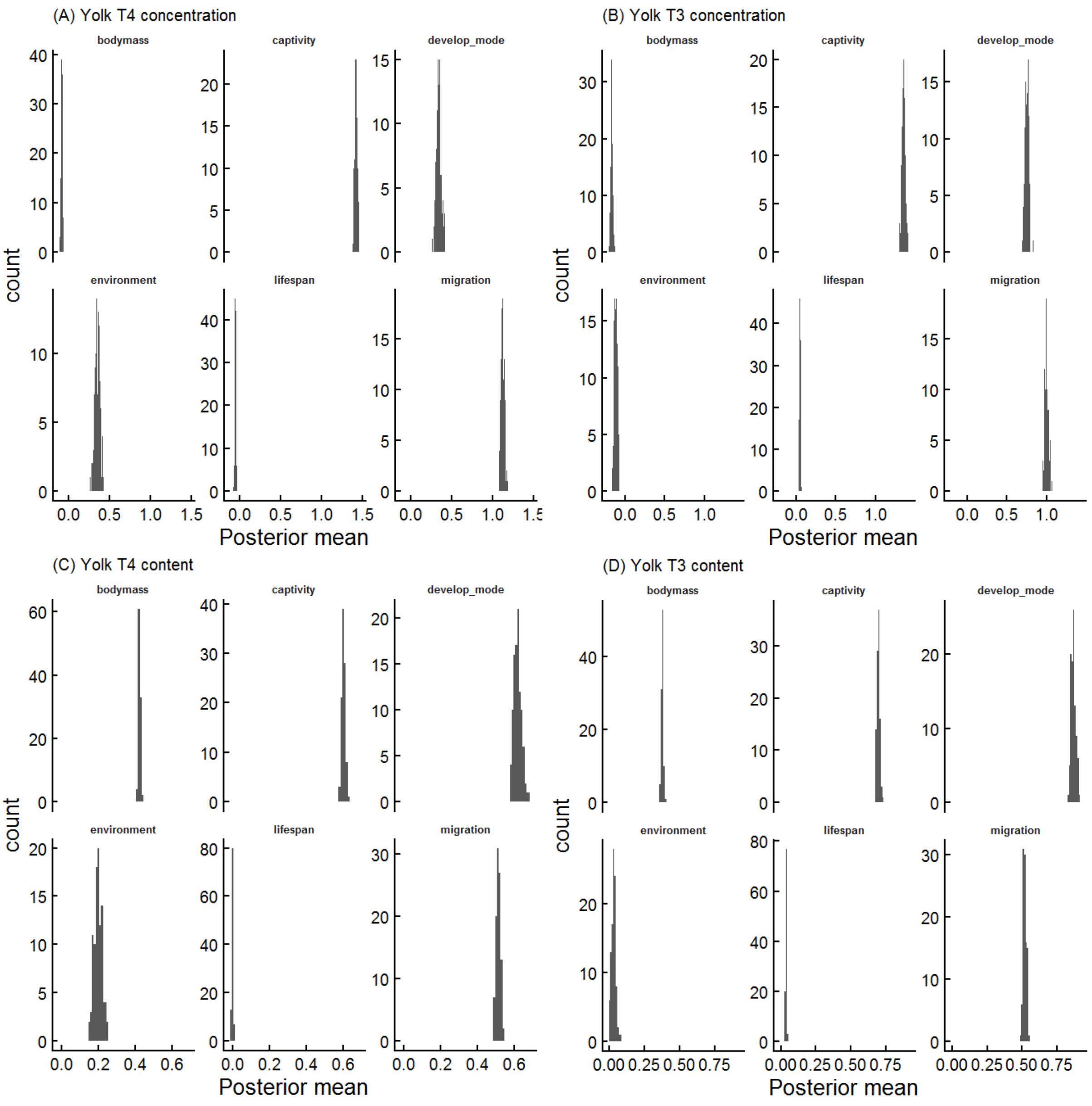


**Figure S1.** Histograms of posterior means for each life-history traits tested in model and data set 1 (see Table 1) across 100 possible phylogenetic trees from the Hackett backbone. All results were highly similar across different trees, indicated by the narrow range of the posterior means.

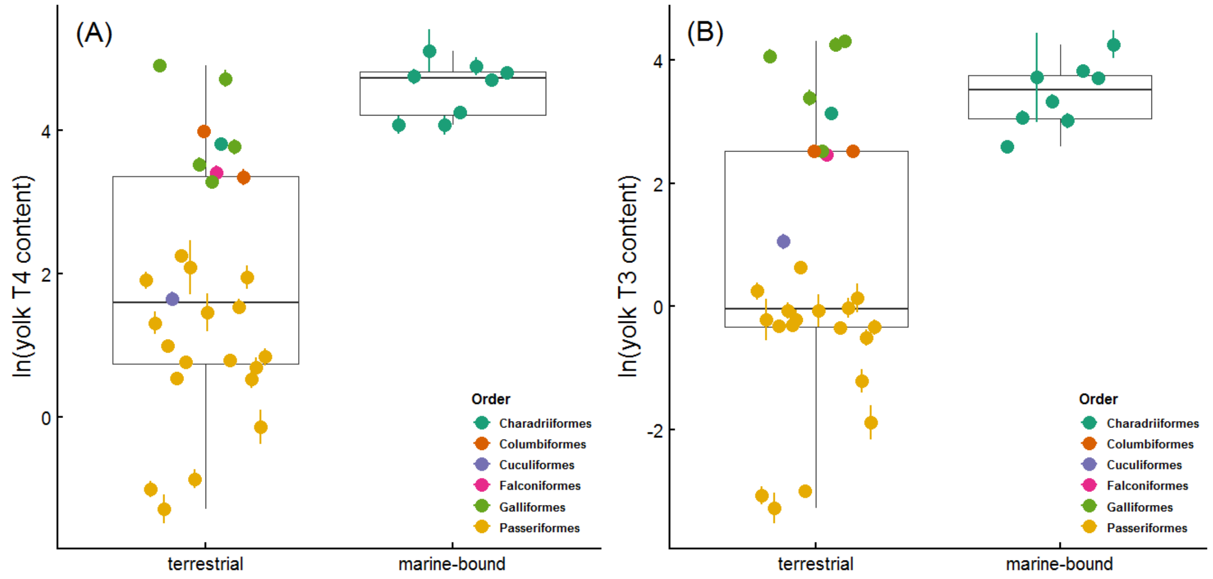


**Figure S2.** Boxplots of yolk T4 (A) and T3 (B) contents (ng/yolk, ln-transformed raw data) between marine-bound and terrestrial species. Boxplots represent the median (the middle line) and the first and the third quartiles (the box), and the whiskers extend to 1.5 times of the interquartile range. Colored dots represent species-specific means (±SE).

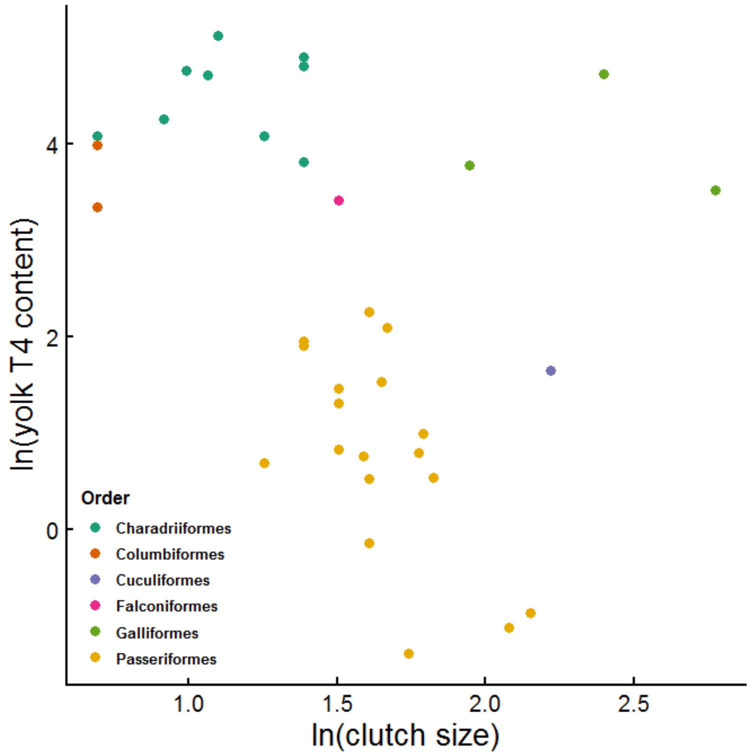


**Figure S3.** Scatterplot of species clutch size and yolk T4 content on a natural-log scale. Dots represent species-specific means and colors represent the orders they belong to.
